# HLA class II mediates type 1 diabetes risk by anti-insulin repertoire selection

**DOI:** 10.1101/2021.09.06.458974

**Authors:** Arcadio Rubio García, Athina Paterou, Mercede Lee, Hubert Sławiński, Ricardo Ferreira, Laurie G Landry, Dominik Trzupek, Luc Teyton, Agnieszka Szypowska, Linda S Wicker, Maki Nakayama, John A Todd, Marcin Ł Pękalski

**Affiliations:** Wellcome Centre for Human Genetics, Nuffield Department of Medicine, University of Oxford, Oxford, United Kingdom; Barbara Davis Center for Childhood Diabetes, University of Colorado School of Medicine, Aurora, Colorado, United States; Department of Immunology and Microbiology, Scripps Research Institute, La Jolla, California, United States; Deparment of Pediatrics, Medical University of Warsaw, Warsaw, Poland

## Abstract

Type 1 diabetes (T1D) is a common autoimmune disorder characterized by the destruction of insulin-secreting pancreatic β cells [1], in which polymorphism of the human leukocyte antigen (HLA) class II region is the major genetic risk factor [2, 3, 4]. However, how variation in class II molecules alters T1D risk remains a longstanding question. Here we show how T1D risk due to HLA class II haplotype combinations [5] correlates with the frequency of negatively charged sequences in the CDR3β region of CD4^+^ T cell receptor (TCR) repertoires purified from peripheral blood. These sequences are known to be common in receptors that bind insulin B:9–23 [6], the primary autoantigen in T1D. We also show the same effect in circulating activated CD4^+^ T cells from newly-diagnosed T1D cases, and in islet-infiltrating T cells from patients with active T1D. Furthermore, we demonstrate that the proportion of insulin-reactive CD4^+^ T cells present in islets is predicted by the frequency of these negatively charged CDR3β amino acid sequences. Our results suggest diagnostic uses of T cell repertoire profiling in early detection of insulin autoimmunity, and inform ongoing efforts to improve tolerance induction to insulin and prevention of T1D [7].

## Main text

The association between genetic variants in the human leukocyte antigen (HLA) region and susceptibility to different infectious, autoimmune and neoplastic disorders emerged more than half a century ago [8]. In case of type 1 diabetes (T1D), this association was initially reported for HLA class I alleles [9, 10]. A stronger genetic signal linked to HLA class II soon followed up [11, 12, 13], with a large portion of the risk mapped to the amino acid at position 57 in the DQB1 molecule [2], in which the presence of aspartic acid (D) correlates with protection against T1D [14].

The T1D associations with HLA class II DR and DQ alleles, individually, or combined in their cis interaction effects on haplotypes and trans effects between haplotypes on different chromosomes are complex [3, 4]. Therefore, here we use class II haplotype combinations in diplotypes to take into account these interactions in our investigation of a possible mechanism underlying their associations with T1D.

HLA variants are known to bias T cell receptor (TCR) repertoires by imposing different interfacing constraints and exposing cells to alternative peptidomes during the development of central and peripheral tolerance [15, 16, 17], which implies repertoire differences may explain the susceptibility or protection conferred by HLA class II haplotypes.

Previously, we assembled a cohort of T1D families, as part of the Diabetes-Genes, Autoimmunity and Prevention Centre (D-GAP), and built a collection of peripheral blood mononuclear cells (PBMCs) from blood samples from children with T1D and their unaffected siblings in order to define the immune system before and during T1D [18]. Using this resource, we investigated possible associations between HLA class II and the TCR repertoire.

### HLA class II haplotypes explain most of the variance in the CDR3 region of CD4^+^ TCR repertoires

We purified circulating CD4^+^ T cells (n = 349623) from D-GAP participants (n = 48, median age = 11 years) with haplotypes across the susceptibility-protection continuum [5]. These cells were stimulated and loaded into a single-cell platform for preparing paired 5’ gene expression and TCR libraries [19]. After sequencing, gene expression data analysis and repertoire assembly, we obtained 288903 cells with both a valid gene expression profile and at least one productive TCR chain (Materials and Methods).

Using CD4^+^ T conventional cells (n = 276581), defined as those outside the regulatory T cell (Treg) cluster, we calculated the frequencies of all 2-mer sequences in the complementarity-determining region 3 (CDR3) α and β grouped by donor. We excluded an N-terminal prefix and a C-terminal suffix from each CDR3 to avoid the HLA binding bias that constrains gene usage [16, 20]. We then considered the subset of donors with susceptible or protected HLA class II haplotypes (Supplementary Tables S2 and S4). We reduced the dimensionality of the resulting matrix (40 donors x 798 2-mers) by solving a multidimensional scaling optimization problem with a given Jaccard distance metric.

Donor repertoires clustered by HLA class II risk group in the resulting 2-dimensional representation of CDR3 sequences (Figure 1), and differences between cluster centroids were significant (*p* = 0.001). This demonstrates that despite age, sex and environmental covariates, which are major factors in the development of T1D, HLA class II haplotypes determine the epitope-binding properties of the CD4^+^ T cell repertoire.

**Figure 1:**
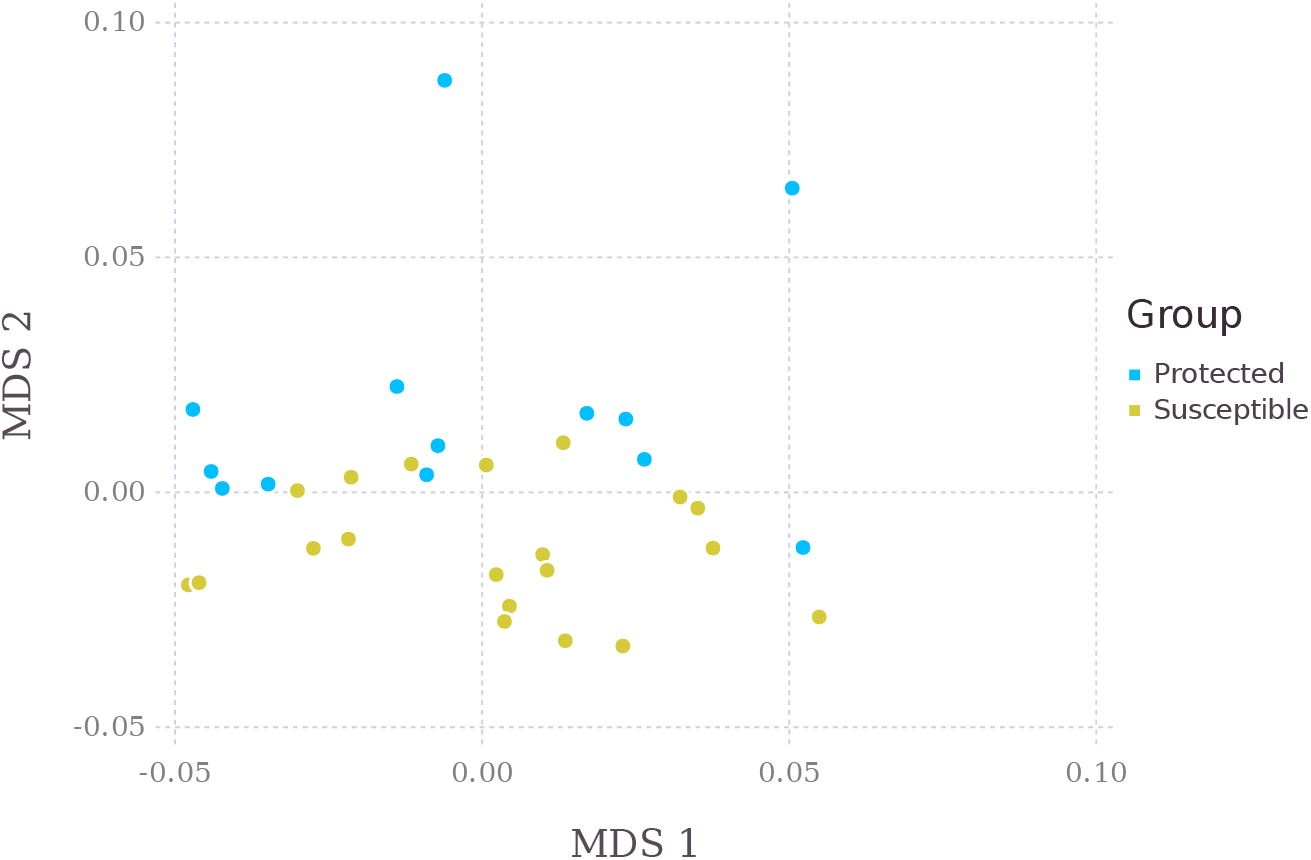
Multidimensional scaling (MDS) of genetically protected and susceptible donor repertoires using combined CDR3α and CDR3β 2-mer frequencies from CD4^+^ conventional T cells.

### CDR3β regions in CD4^+^ TCRs from genetically susceptible donors exhibit a higher frequency of negatively charged residues

Next, we examined which specific differences on CDR3 selection pressure could be explained by HLA class II risk using a multilevel model (Materials and Methods). This model estimates differences in frequencies as a function of T1D risk due to HLA class II haplotype interactions from the observed counts of each CDR3 k-mer, as well as differences and fold changes across both extremes of the disease risk spectrum.

Irrespective of the selected k-mer size, we found sequences with one or more negatively-charged residues (D or glutamic acid, E) were more likely to be present in β chains belonging to donors who carried a susceptible HLA class II haplotype (Figures 2 and 3). For the special case k = 1, which reduces to single amino acid frequencies, we used a linear model to estimate differences. Sorting CDR3β amino acids by their average interaction free energy [21] revealed negatively charged residues D and E are often replaced by high interaction potential ones (leucine L, isoleucine I and valine V; Figure S1) as class II-associated disease risk decreases. False discovery rates (FDR) for the regression slopes of D (FDR = 8e-9) and V (FDR = 1.45e-3) provide additional support for non-zero effects (Figure 4).

**Figure 2:**
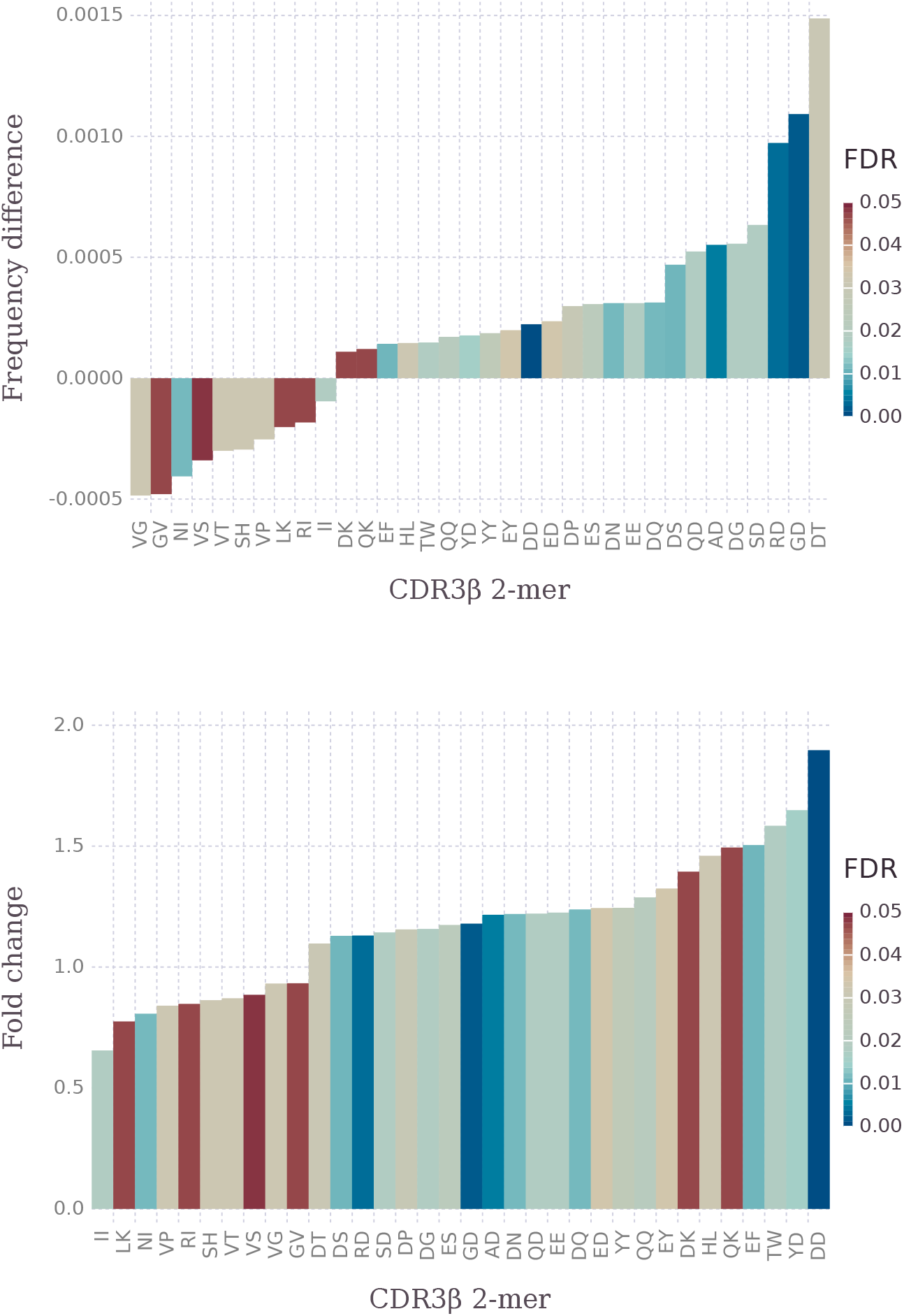
Estimated CDR3β 2-mer frequency differences and fold changes across HLA class II risk extremes (Table S1) in CD4^+^ conventional T cells.

**Figure 3:**
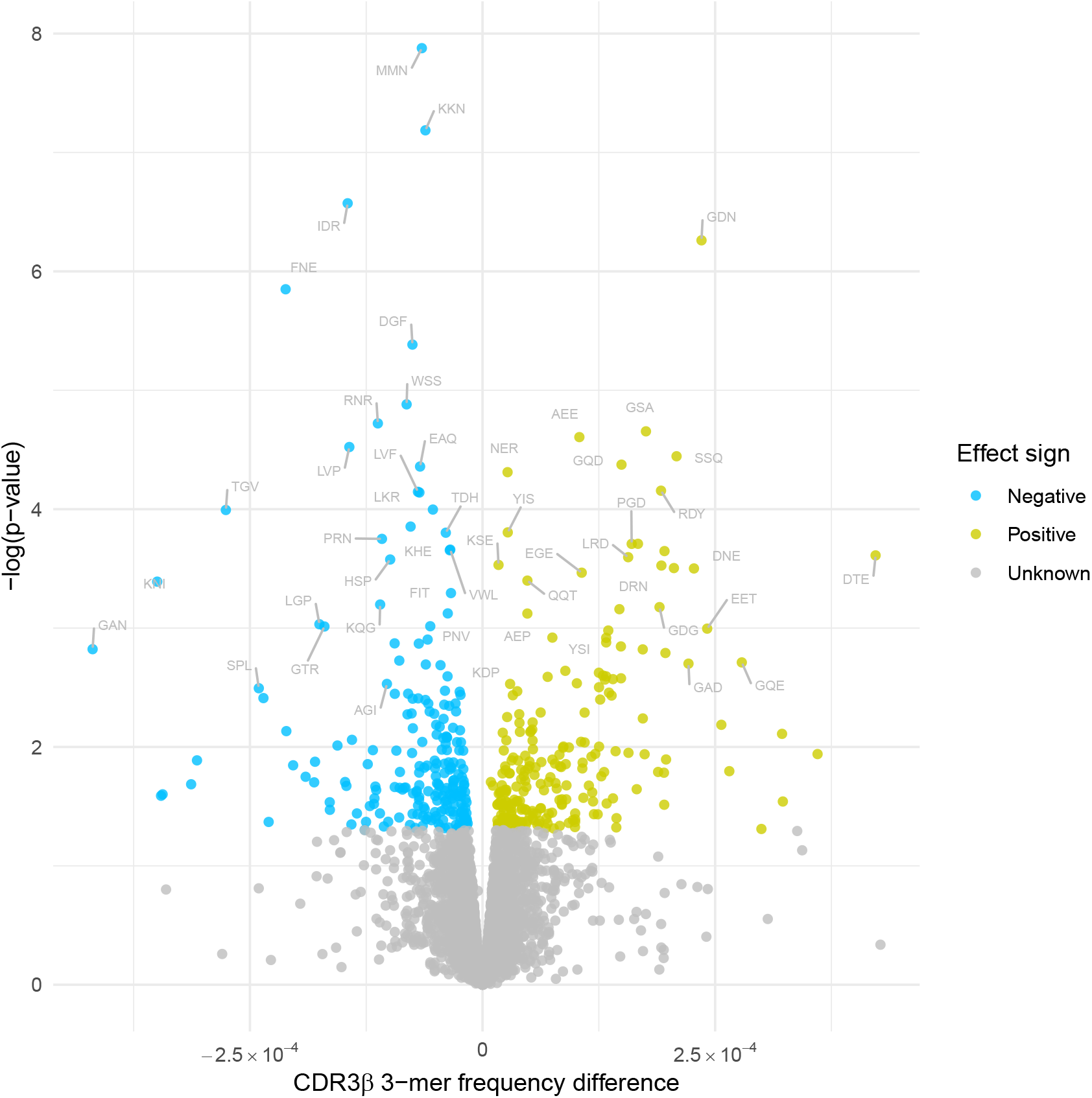
Estimated CDR3β 3-mer frequency differences across HLA class II risk extremes (Table S1) in CD4^+^ conventional T cells.

**Figure 4:**
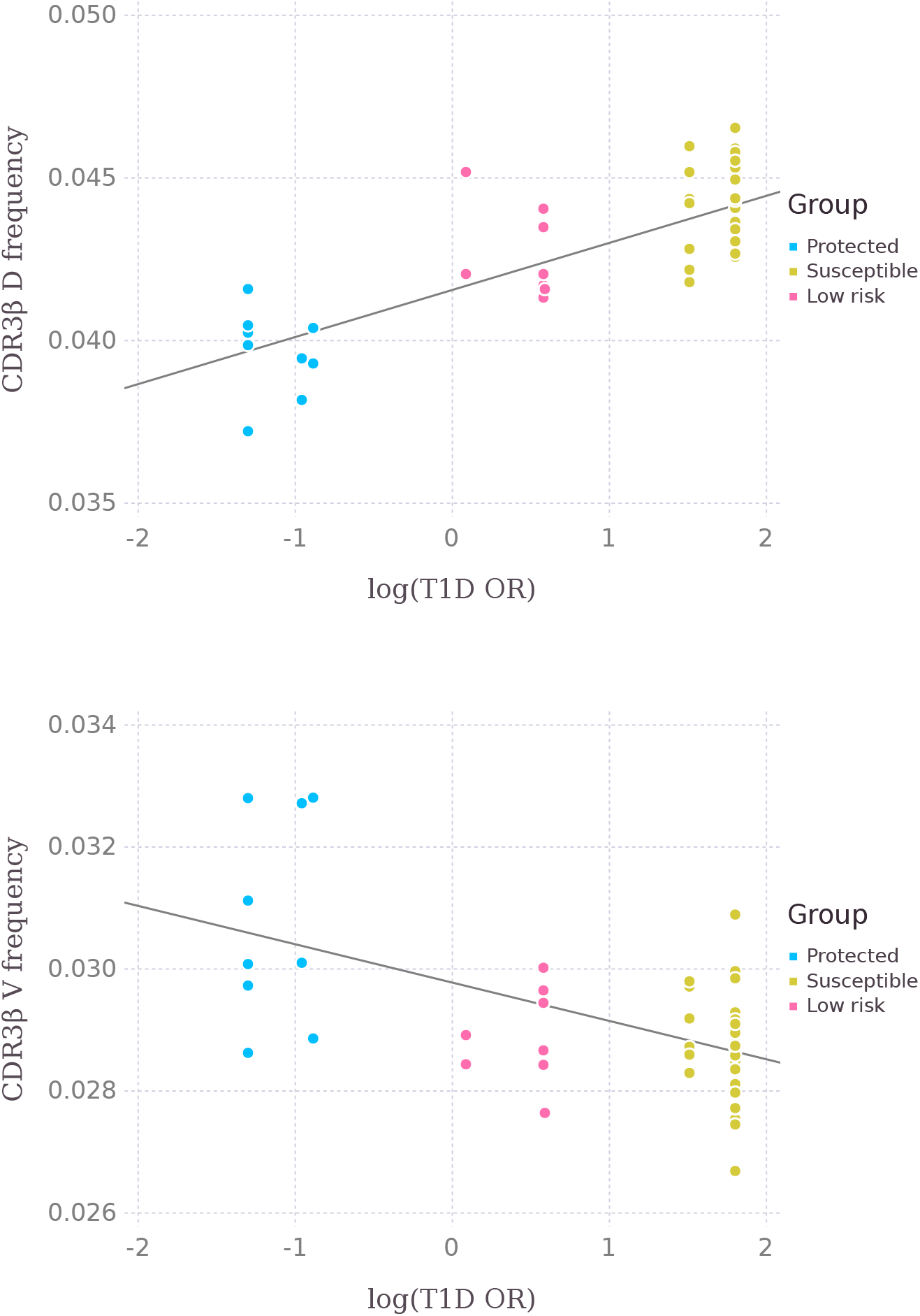
Observed CDR3β amino acid frequencies for aspartic acid (D) and valine (V) in CD4^+^ conventional T cells regressed against T1D log odds risk due to HLA class II interactions (Table S1).

The DQB1 57 association with T1D is conserved between humans and mice [2]. The non-obese diabetic (NOD) mouse develops autoimmune diabetes spontaneously and has S 57 in the β chain of the ortholog of DQ molecule, I-A^g7^, compared to autoimmune diabetes-resistant strains that have D 57. Our TCR results in humans show a remarkable evolutionary conservation with the previous class II-TCR findings in NOD mice, in which mutation of I-A^g7^ chain at position 57 from S to D resulted in T1D protection, and reduced D and E amino acids in the CDR3 sequences of insulin-specific CD4^+^ T cells in the mutant strain compared to the wild-type NOD mice [6].

In case of Tregs, we observed the same trend as in conventional T cells (Figure 5), which suggests this amino acid bias occurs during positive thymic selection. It is also possible that the repertoires of Tregs induced during the development of peripheral tolerance are affected by HLA class II polymorphisms [22].

**Figure 5:**
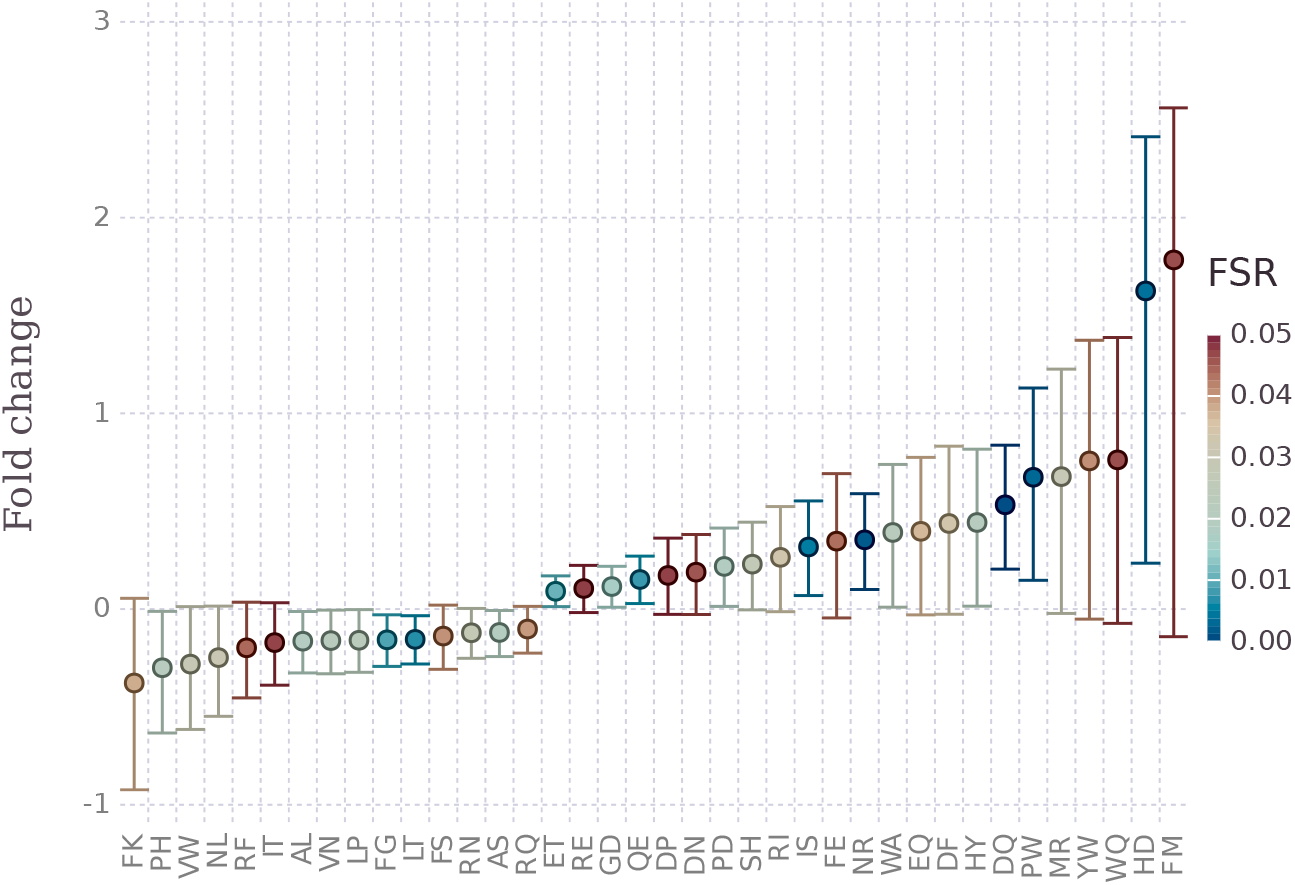
Estimated CDR3β 2-mer log fold changes across HLA class II risk extremes (Table S1) in CD4^+^ regulatory T cells.

In case of α chains, we also estimated systematic differences at low false discovery rates (Figure S3). These seem to correspond to biases in the usage of particular Vα and Jα genes, which are carried onto the CDR3α region due to its low recombination diversity. In particular, top estimated differences in k-mers match fragments of `SGTYK`, which is a sequence encoded in the TRAJ40 gene whose usage is upregulated in individuals who carry susceptible HLA class II haplotypes.

### HLA class II susceptible donors are more likely to select CD4^+^ TCRs that recognize insulin B:9–23

We then investigated whether these observed repertoire differences correlate with an immune response against the primary epitope in T1D, insulin B:9–23 [23, 24, 25]. We sorted activated circulating CD4^+^ T cells (n = 19969), defined as HLA-DR^+^ CD38^+^, from a cohort of newly diagnosed children (n = 8) carrying susceptible HLA class II haplotypes. We also sourced CD4^+^ T cell receptor clonotypes (n = 1428) isolated from the islets of individuals (n = 5) in the Network of Pancreatic Organ Donors (nPOD) [26] who had active T1D and also carried susceptible haplotypes. We compared these against all the individuals with maximum susceptibility (n = 23) in our original cohort (Supplementary Table S1), and therefore a D frequency distribution with the highest mean (Figure 4). Both CDR3β sequences from islets and circulating activated cells had a significantly higher frequency of aspartic acid (*p* = 0.0097 and 0.0034, respectively; Figure 6).

**Figure 6:**
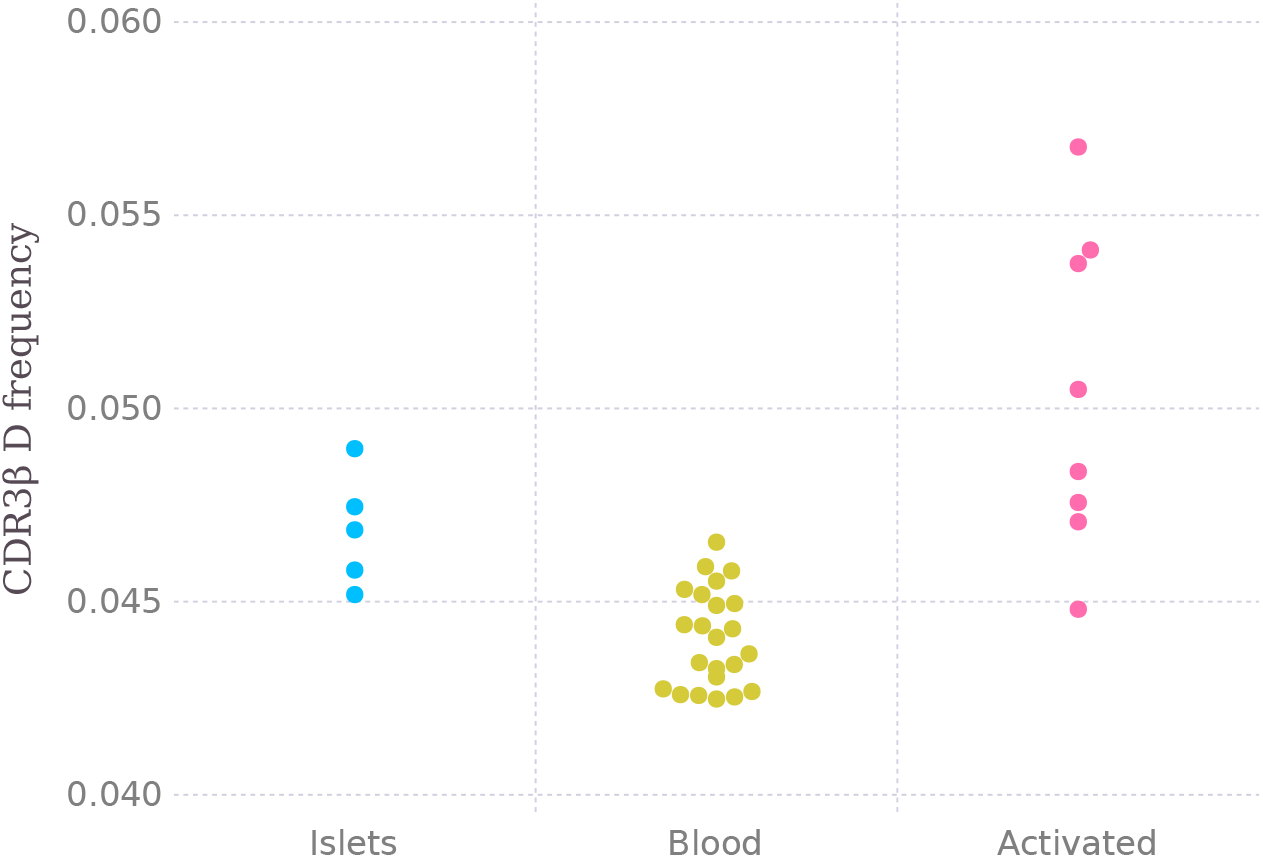
CDR3β aspartic acid (D) frequency measured in CD4^+^ T cells infiltrating islets, circulating CD4^+^ T cells from susceptible HLA class II donors and activated circulating CD4^+^ T cells from recently diagnosed T1D donors.

Finally, we analyzed clonotypes (n = 159) from nPOD donors (n = 5) where reactivity against preproinsulin had been tested [27] to validate our hypothesis that negative charges correlate with the presence of insulin-reactive TCRs. The slope of a linear model provided strong evidence of an association between both variables (*p* = 0.0008). Alternatively, a binomial regression model was also consistent with these results (Figure 7; *p* = 0.0843).

**Figure 7:**
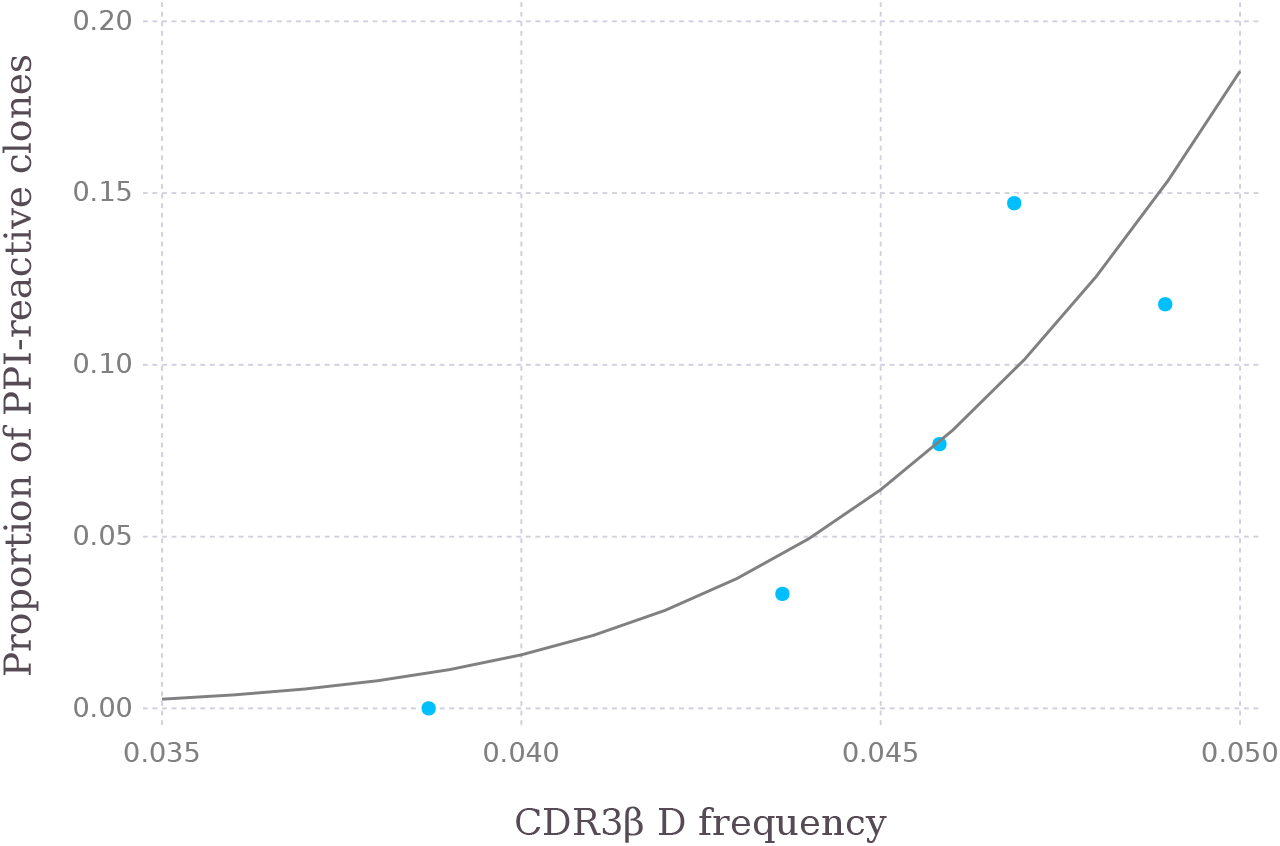
Proportion of preproinsulin-reactive CD4^+^ T cell clones present in islets as a function of CDR3β aspartic acid (D) frequency in CD4^+^ T cells infiltrating islets.

## Discussion

Autoimmune diseases have long been hypothesized to be, essentially, diseases of the receptor repertoire [28, 29]. Large observational studies employing bulk RNA and TCR sequencing have uncovered broad associations such as gene usage biases [16] or pathogen-specific receptor sequences [17] linked to certain HLA alleles. Disease-specific efforts related to T1D have not been able to include this fundamental covariate in their experimental design, and have mostly drawn inconclusive results [30, 31, 32].

Our study offers a mechanistic explanation of T1D risk due to HLA class II haplotypes in terms of CDR3β sequence biases. Susceptibility to T1D correlates with the frequency of negatively-charged sequences in the peptide epitope-binding region of TCR β chains. These negative charges are common in receptors that bind insulin B:9–23, the primary autoantigen in T1D, as demonstrated in a recent mouse model [6]. Thus, higher genetic susceptibility is associated with a larger proportion of insulin-reactive CD4^+^ T cells. As predicted by the Miyazawa-Jerningan contact energies model [21], favorable interactions with a D residue are those occurring with phenylalanine, isoleucine and valine, which match the sequences we observed to be enriched in class II-protected individuals, and depleted from susceptible ones.

Understanding what aspects of TCR repertoires play a role in the onset of autoimmunity could contribute to the development of better biomarkers and effective immunotherapies. The inclusion of biomarkers derived from immune repertoires, which capture risk from both environmental factors [33] and their interactions with host genetics, might add to the predictive value of T1D genetic risk scores and autoantibody testing.

A fundamental and related open question is to determine what events precipitate insulin autoimmunity. Functional repertoire differences observed in susceptible HLA backgrounds can also influence recognition of insulin B:9–23 epitope mimics expressed in the microbiome. We have previously demonstrated the presence of insulin B:9–23 mimotopes within the transketolase (TKT) superfamily of enzymes, which are highly upregulated during infant weaning, at the same age as the peak of anti-insulin autoimmunity [34]. Notwithstanding the peripheral presentation of TKT to T cells, TKT-expressing microbes can also be transported into the thymus by the subset of specialized dendritic cells causing dysbiosis-dependent development of TKT-specific autoreactive T cells [35]. These considerations warrant further investigation into the immunological and phenotypic effects of bacterial probiotic supplementation at the molecular level in the context of primary prevention of autoimmunity and allergy, as well as host genetic susceptibility and resistance.

## Acknowledgments

We thank all donors and patients participating in this study. We are grateful to the study volunteers and staff associated with D-GAP, including the Wellcome Clinical Research Facility (Addenbrooke’s Clinical Research Centre, Cambridge, UK) and DMech hospital sites, including the Institute of Mother and Child (Warsaw, Poland). We are grateful to Wojciech Szypowski from the Polish Society for Autoimmune Diseases (Warsaw, Poland) for sample logistics. DMech blood donors were recruited under the South Central Oxford A Ethics Committee (Ethics Reference: 18/SC/0559, IRAS Project ID: 243305). We would like to thank members from the Diabetes and Inflammation Laboratory (University of Oxford, Oxford, UK) Heather McMurray, Shannah Donhou, Sarune Kacinskaite, Michael Ellis, Sandra Banks, Georgina Burton and past members at the University of Cambridge (Cambridge, UK) led by Helen Stevens for help with blood sample processing. We would also like to thank Claire Scudder, Raqeem Mahmood and Sylwia Kopjasz for managing ethics and blood donor recruitment; Hong Harper for managing the patients’ registry and D-GAP data; Florent Yvon for HLA imputation of D-GAP patients; Jamie Inshaw for information concerning D-GAP donor metadata; and Olga Platonova for managing finance and funding. We would like to thank Moustafa Attar, Oxford Genomics Centre (University of Oxford), for assistance with single cell sequencing. We would like to dedicate this work to the memory of David B Dunger (1948–2021).

## Materials and methods

### Donor selection

Donors (Table S1) were obtained from the Diabetes Genes, Autoimmunity and Prevention (D-GAP) cohort which comprised T1D cases and unaffected siblings (REC Ref: 08/H072025) [18]. This cohort provided PBMCs and genomic DNA samples. Genomic DNA was prepared from PBMCs or whole blood using QiaAmp DNA Blood kit (Qiagen), or phenol/chloroform extraction.

Initially, we selected two groups with an equal number of donors in the two extremes of the T1D susceptibility-protection axis, namely DR3/4 or susceptible (Table S2) and DR15 or protected (Table S4). This choice was performed using data from Taqman sequencing (Applied Biosystems) of four SNPs (rs2187668, rs660895, rs9271366 and rs7454108) and RELI SSO (DYNAL Biotech) classical HLA typing.

Due to the limited number of DR15 homozygotes, we also included DR15 heterozygotes with a neutral haplotype—a residue other than the susceptible Ala (A) at DQB1 position 57. For example, some of those DR15 heterozygotes had protective Asp (D) along with a neutral Val (V) or Ser (S).

In each batch loaded on a single-cell Chromium V(D)J cassette (10x Genomics), wherever possible, we matched individuals for age (< 20 years old) and homozygosity for DQB1 position 57. For example, in each batch all susceptible DR3/4 individuals were homozygotes for DQB1 Ala (A) 57, and the protected DR15 were an equal proportion of homozygotes and heterozygotes of DQB1 Asp (D) 57.

We also included two batches of DR3/DR4 T1D patients and two batches of DR4/DQ8 vs DR4/DQ7 or low risk (Table S3) homozygotes controlled focusing on DQB1 position 57. HLA types were further confirmed with ImmunoArray-24 BeadChip v2.0 (Infinium) or HumanImmuno BeadChip v1.0 (Illumina) and HLA imputation [36], along with further SSP classical HLA typing (MC Diagnostics and Oxford Transplant Centre).

### Next-generation sequencing

We washed CD4^+^ T cells in PBS with 0.04% BSA and re-suspended them at a concentration of ~800–1200 cells/μl, before capturing single cells in droplets using the Chromium platform (10x Genomics). Generation of paired gene expression and T-cell receptor libraries was performed using the Chromium Single Cell V(D)J Reagent Kits v1 and v1.1b. We quantified cDNA using Qubit dsDNA HS Assay Kit (Life Technologies) and High Sensitivity D5000 ScreenTape (Agilent). Quantification of libraries was carried out using Qubit dsDNA HS Assay Kit (Life Technologies) and D1000 ScreenTape (Agilent).

Libraries were sequenced on HiSeq 4000 and NovaSeq 6000 (Illumina) to achieve an average of 20000 reads per cell for gene expression libraries and 5000 read pairs per cell for TCR libraries.

### Single-cell data processing

We preprocessed all single cell RNA and TCR sequence libraries separately using Cell Ranger v4.0.0 (10x Genomics) to obtain gene counts and receptor assemblies for each donor. These were then merged into a single gene expression matrix and a single TCR database, which mapped gene counts or TCR chains to cells and donors.

Subsequently, we called cells from read counts with a minimum of 300 genes expressed. We also removed genes not present in at least 50 cells to keep the expression matrix tractable. Furthermore, we applied batch-dependent cutoffs to remove outliers suspected to be cell doublets or multiplets. We also filtered cells with more than 15% of mitochondrial expression to discard those undergoing apoptosis. After data cleanup, we normalized all expression values to 10^4^ reads per cell and applied a logarithmic transformation. Next, we discarded all but the top 5000 most variable genes, and regressed out differences due to sequencing depth and mitochondrial expression.

Lastly, we aligned cells from each sample using batch-balanced nearest neighbors [37], reduced the dimensionality [38], called clusters [39], and performed a multivariate differential expression [40] to find population markers. The initial run yielded two low-frequency clusters with non-CD4^+^ contaminants. We discarded cells mapping to these, reran all data processing steps, found another cluster with contaminants, and iterated through the same process one last time to remove another non-CD4^+^ cluster. This lead to 12 different cell subpopulations (Figures S6 and S7), with a distinct FOXP3^+^ CTLA4^+^ regulatory T cell cluster (Figure S8).

We filtered TCRs called by Cell Ranger to retain consensus assemblies with productive rearrangements only. Finally, we performed an inner join between gene expression and receptor assembly data using cell barcodes to obtain TCR chains paired with gene expression cluster information.

### Estimation of TCR repertoire differences

We used an overdispersed beta-binomial model (Equation 1) to regress k-mer counts as a function of T1D risk due to HLA class II haplotypes, while accounting for a significant variability in the number of observed TCR chains per donor and in the observed frequencies across donors with similar risk.

The model assumes that counts observed for each k-mer and each donor *k_ij_* follow a binomial distribution, which has been sampled from a latent donor-specific beta random variable *θ_ij_*. In turn, all donor-specific betas are generated by a linear model whose output is transformed by a complementary log-log inverse link function. The particular objective of inference is to estimate the latent probability *μ_i_*, as well as the derived variables that represent difference and fold changes, *δ_i_* and *ρ_i_*, which are calculated using susceptible and protective risk extremes (Table S1).

It is important to note we used a reparametrization of beta distributions in terms of mean and sample size or pseudo-dispersion, which makes the generative process easier to express.

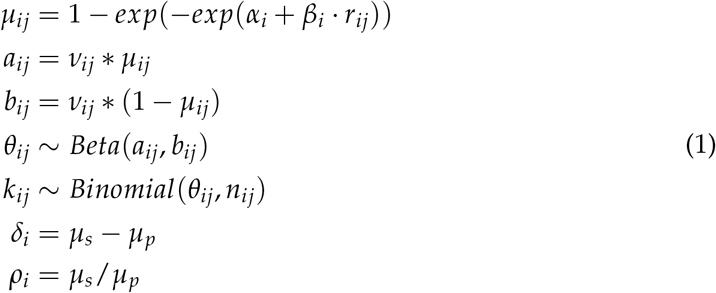

We initialized 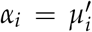, the generation probability of each k-mer estimated by a thymic recombination model [41], and maximized the likelihood implicitly defined by the generative process described above. p-values were derived from a likelihood ratio test against a null model defined as a linear model without a slope term.

## Supplementary material

**Table S1:** Sample selection and HLA class II haplotype interactions. T1D odds ratios (OR) for DQ-DR haplotype interactions were obtained from UK Biobank [5]. Estimates for the maximum (63.17) and minimum (0.05) OR are used to calculate differences and fold changes, with estimates for the maximum OR serving as baseline (Materials and Methods).

| n | Haplotype 1 |  | Haplotype 2 |  | DQB1<sub>57</sub> | OR | Group |
| --- | --- | --- | --- | --- | --- | --- | --- |
|  | DQA1 | DQB1 | DQA1 | DQB1 |  |  |  |
| 23 | 03:01 | 03:02 | 05:01 | 02:01 | AA | 63.17 | Susceptible |
| 8 | 03:01 | 03:02 | 03:01 | 03:02 | AA | 32.30 | Susceptible |
| 1 | 03:01 | 03:01 | 06:01 | 03:01 | DD | 3.88 | Low risk |
| 5 | 03:02 | 03:01 | 03:02 | 03:01 | DD | 3.80 | Low risk |
| 1 | 03:03 | 03:01 | 05:05 | 03:01 | DD | 1.22 | Low risk |
| 1 | 03:01 | 03:01 | 05:05 | 03:01 | DD | 1.22 | Low risk |
| 2 | 01:02 | 06:02 | 05:05 | 03:01 | DD | 0.13 | Protected |
| 2 | 01:02 | 06:02 | 01:02 | 06:02 | DD | 0.11 | Protected |
| 2 | 01:02 | 06:02 | 05:05 | 06:03 | DD | 0.05 | Protected |
| 2 | 01:02 | 06:02 | 01:02 | 05:02 | DS | 0.05 | Protected |
| 1 | 01:02 | 06:02 | 01:02 | 06:04 | DV | 0.05 | Protected |

**Table S2:**
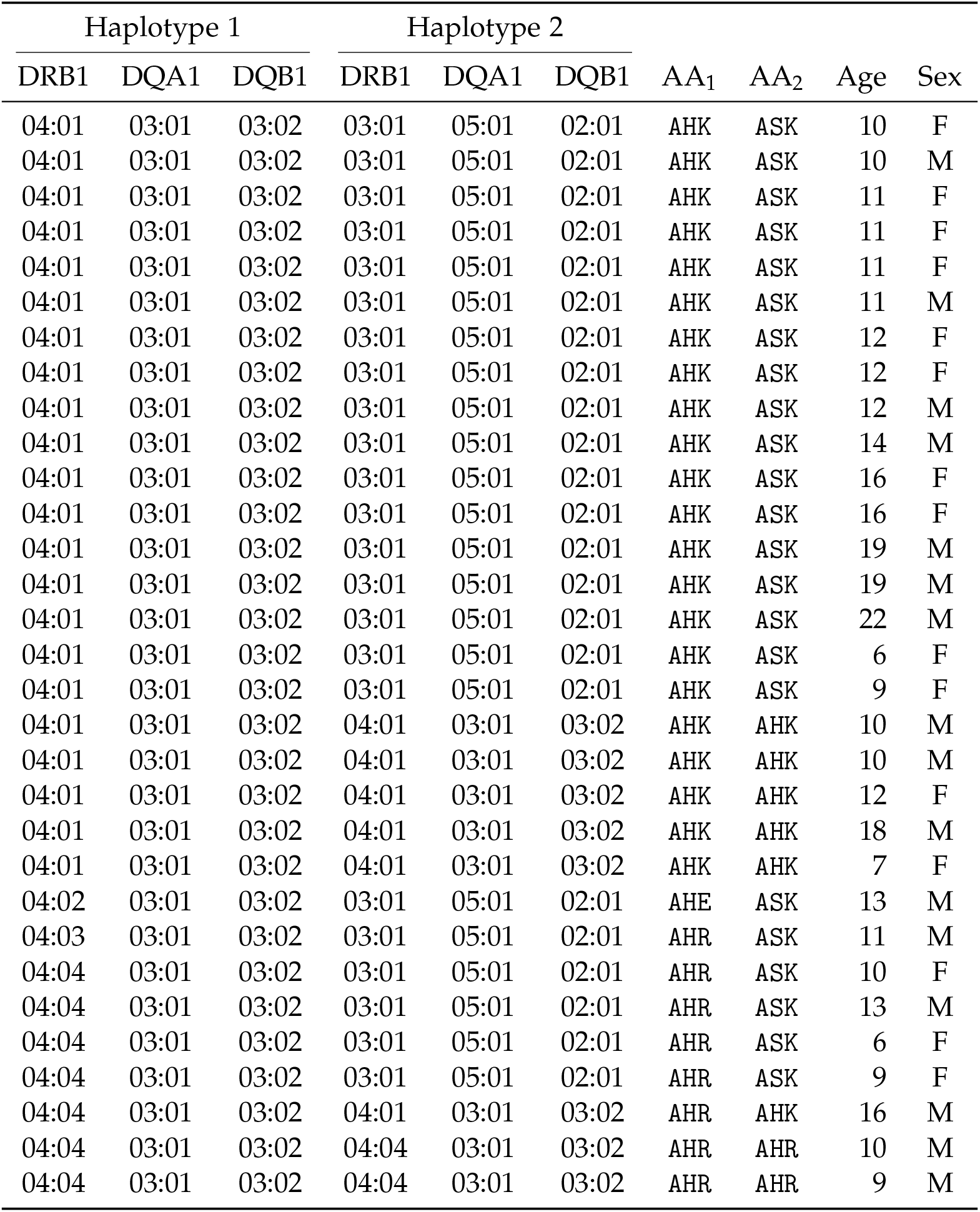
Genetically susceptible group. The strings AA_1_ and AA_2_ encode amino acids at HLA DQB1 57, DRB1 13 and 71 [3], respectively.

| Haplotype 1 |  |  | Haplotype 2 |  |  | AA<sub>1</sub> | AA<sub>2</sub> | Age | Sex |
| --- | --- | --- | --- | --- | --- | --- | --- | --- | --- |
| DRB1 | DQA1 | DQB1 | DRB1 | DQA1 | DQB1 |  |  |  |  |
| 04:01 | 03:01 | 03:02 | 03:01 | 05:01 | 02:01 | AHK | ASK | 10 | F |
| 04:01 | 03:01 | 03:02 | 03:01 | 05:01 | 02:01 | AHK | ASK | 10 | M |
| 04:01 | 03:01 | 03:02 | 03:01 | 05:01 | 02:01 | AHK | ASK | 11 | F |
| 04:01 | 03:01 | 03:02 | 03:01 | 05:01 | 02:01 | AHK | ASK | 11 | F |
| 04:01 | 03:01 | 03:02 | 03:01 | 05:01 | 02:01 | AHK | ASK | 11 | F |
| 04:01 | 03:01 | 03:02 | 03:01 | 05:01 | 02:01 | AHK | ASK | 11 | M |
| 04:01 | 03:01 | 03:02 | 03:01 | 05:01 | 02:01 | AHK | ASK | 12 | F |
| 04:01 | 03:01 | 03:02 | 03:01 | 05:01 | 02:01 | AHK | ASK | 12 | F |
| 04:01 | 03:01 | 03:02 | 03:01 | 05:01 | 02:01 | AHK | ASK | 12 | M |
| 04:01 | 03:01 | 03:02 | 03:01 | 05:01 | 02:01 | AHK | ASK | 14 | M |
| 04:01 | 03:01 | 03:02 | 03:01 | 05:01 | 02:01 | AHK | ASK | 16 | F |
| 04:01 | 03:01 | 03:02 | 03:01 | 05:01 | 02:01 | AHK | ASK | 16 | F |
| 04:01 | 03:01 | 03:02 | 03:01 | 05:01 | 02:01 | AHK | ASK | 19 | M |
| 04:01 | 03:01 | 03:02 | 03:01 | 05:01 | 02:01 | AHK | ASK | 19 | M |
| 04:01 | 03:01 | 03:02 | 03:01 | 05:01 | 02:01 | AHK | ASK | 22 | M |
| 04:01 | 03:01 | 03:02 | 03:01 | 05:01 | 02:01 | AHK | ASK | 6 | F |
| 04:01 | 03:01 | 03:02 | 03:01 | 05:01 | 02:01 | AHK | ASK | 9 | F |
| 04:01 | 03:01 | 03:02 | 04:01 | 03:01 | 03:02 | AHK | AHK | 10 | M |
| 04:01 | 03:01 | 03:02 | 04:01 | 03:01 | 03:02 | AHK | AHK | 10 | M |
| 04:01 | 03:01 | 03:02 | 04:01 | 03:01 | 03:02 | AHK | AHK | 12 | F |
| 04:01 | 03:01 | 03:02 | 04:01 | 03:01 | 03:02 | AHK | AHK | 18 | M |
| 04:01 | 03:01 | 03:02 | 04:01 | 03:01 | 03:02 | AHK | AHK | 7 | F |
| 04:02 | 03:01 | 03:02 | 03:01 | 05:01 | 02:01 | AHE | ASK | 13 | M |
| 04:03 | 03:01 | 03:02 | 03:01 | 05:01 | 02:01 | AHR | ASK | 11 | M |
| 04:04 | 03:01 | 03:02 | 03:01 | 05:01 | 02:01 | AHR | ASK | 10 | F |
| 04:04 | 03:01 | 03:02 | 03:01 | 05:01 | 02:01 | AHR | ASK | 13 | M |
| 04:04 | 03:01 | 03:02 | 03:01 | 05:01 | 02:01 | AHR | ASK | 6 | F |
| 04:04 | 03:01 | 03:02 | 03:01 | 05:01 | 02:01 | AHR | ASK | 9 | F |
| 04:04 | 03:01 | 03:02 | 04:01 | 03:01 | 03:02 | AHR | AHK | 16 | M |
| 04:04 | 03:01 | 03:02 | 04:04 | 03:01 | 03:02 | AHR | AHR | 10 | M |
| 04:04 | 03:01 | 03:02 | 04:04 | 03:01 | 03:02 | AHR | AHR | 9 | M |

**Table S3:** Low genetic risk group. The strings AA_1_ and AA_2_ encode amino acids at HLA DQB1 57, DRB1 13 and 71 [3], respectively. The DQA1 03:02 haplotype is actually unresolved to 03:02 or 03:03.

| Haplotype 1 |  |  | Haplotype 2 |  |  | AA<sub>1</sub> | AA<sub>2</sub> | Age | Sex |
| --- | --- | --- | --- | --- | --- | --- | --- | --- | --- |
| DRB1 | DQA1 | DQB1 | DRB1 | DQA1 | DQB1 |  |  |  |  |
| 04:01 | 03:01 | 03:01 | 11:03 | 05:05 | 03:01 | DHK | DSE | 20 | M |
| 04:01 | 03:01 | 03:01 | 12:02 | 06:01 | 03:01 | DHK | DGR | 17 | F |
| 04:01 | 03:02 | 03:01 | 04:01 | 03:02 | 03:01 | DHK | DHK | 11 | M |
| 04:01 | 03:02 | 03:01 | 04:01 | 03:02 | 03:01 | DHK | DHK | 5 | M |
| 04:01 | 03:02 | 03:01 | 04:01 | 03:02 | 03:01 | DHK | DHK | 8 | M |
| 04:01 | 03:02 | 03:01 | 04:01 | 03:02 | 03:01 | DHK | DHK | 8 | M |
| 04:01 | 03:02 | 03:01 | 04:01 | 03:02 | 03:01 | DHK | DHK | 9 | F |
| 04:07 | 03:03 | 03:01 | 01:03 | 05:05 | 03:01 | DHR | DFE | 10 | M |

**Table S4:** Genetically protected group. The strings AA_1_ and AA_2_ encode amino acids at HLA DQB1 57, DRB1 13 and 71 [3], respectively.

| Haplotype 1 |  |  | Haplotype 2 |  |  | AA<sub>1</sub> | AA<sub>2</sub> | Age | Sex |
| --- | --- | --- | --- | --- | --- | --- | --- | --- | --- |
| DRB1 | DQA1 | DQB1 | DRB1 | DQA1 | DQB1 |  |  |  |  |
| 15:01 | 01:02 | 06:02 | 11:04 | 05:05 | 03:01 | DRA | DSR | 20 | M |
| 15:01 | 01:02 | 06:02 | 11:04 | 05:05 | 06:03 | DRA | DSR | 18 | M |
| 15:01 | 01:02 | 06:02 | 11:04 | 05:05 | 06:03 | DRA | DSR | 19 | F |
| 15:01 | 01:02 | 06:02 | 12:01 | 05:05 | 03:01 | DRA | DGR | 18 | F |
| 15:01 | 01:02 | 06:02 | 13:02 | 01:02 | 06:04 | DRA | VSE | 19 | F |
| 15:01 | 01:02 | 06:02 | 15:01 | 01:02 | 06:02 | DRA | DRA | 6 | M |
| 15:01 | 01:02 | 06:02 | 15:01 | 01:02 | 06:02 | DRA | DRA | 7 | M |
| 15:01 | 01:02 | 06:02 | 16:01 | 01:02 | 05:02 | DRA | SRR | 12 | M |
| 15:01 | 01:02 | 06:02 | 16:01 | 01:02 | 05:02 | DRA | SRR | 15 | F |

**Figure S1:**
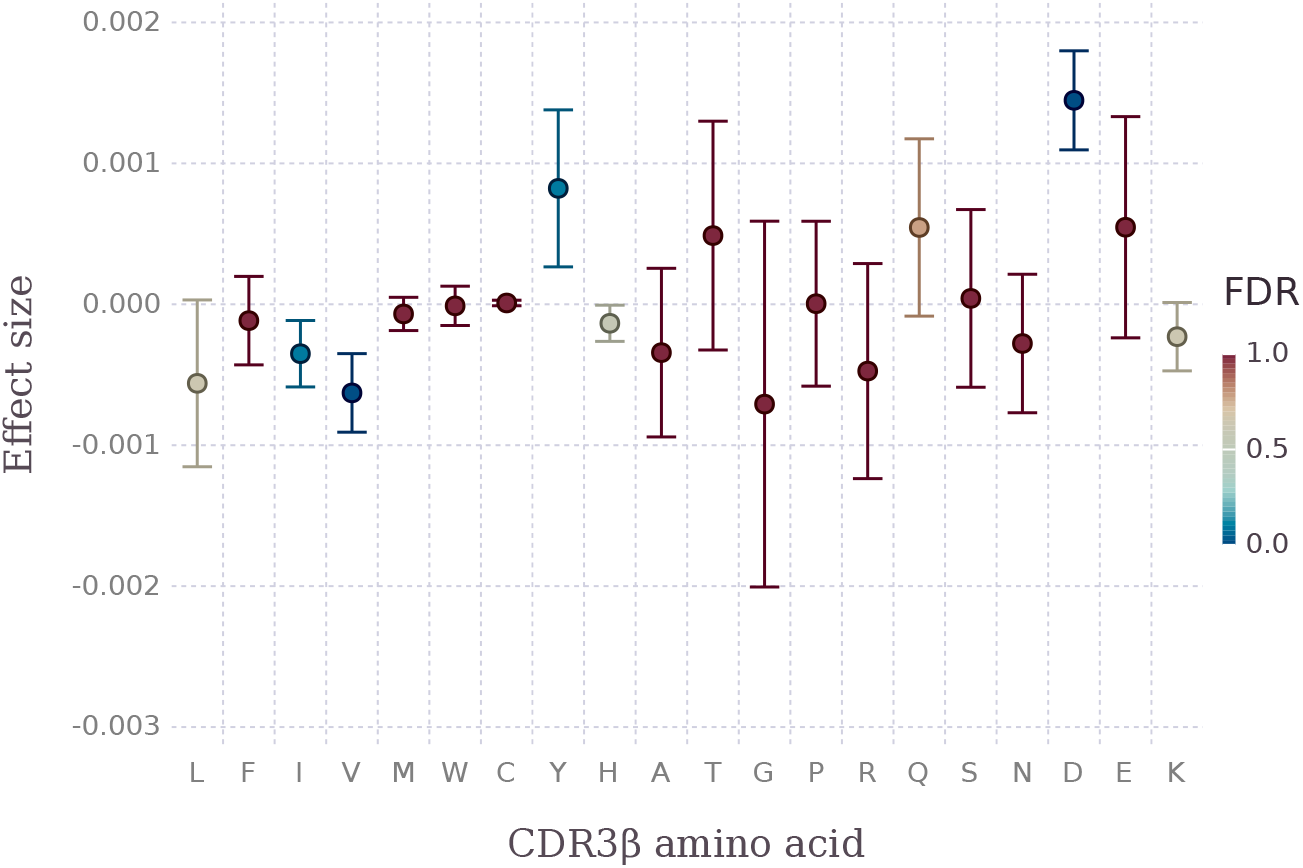
Estimated effects on CDR3β amino acid frequencies from CD4^+^ conventional T cells explained by HLA class II log odds risk. Effects correspond to changes in amino acid frequency with an increase in one log(OR) unit.

**Figure S2:**
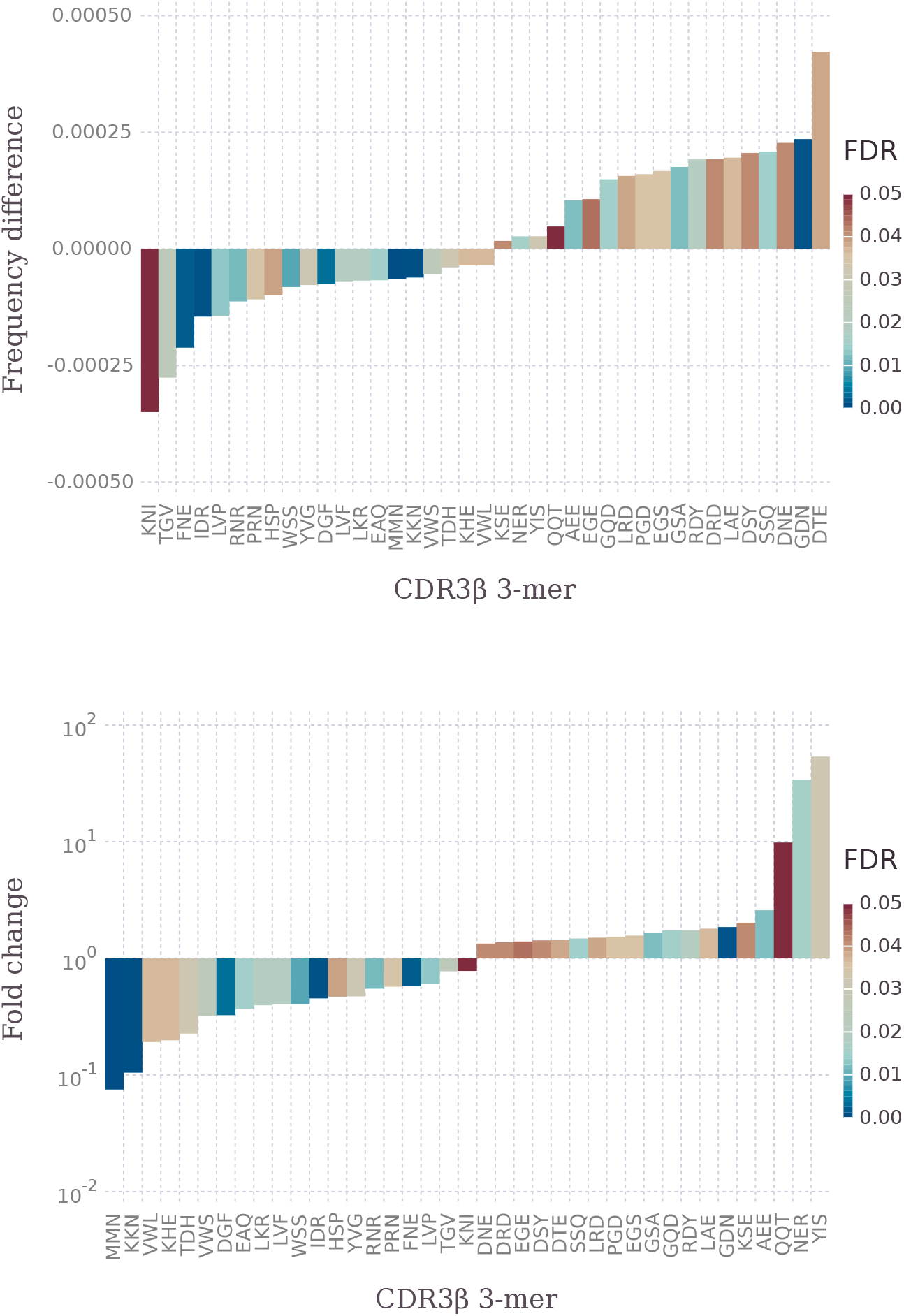
Estimated CDR3β 3-mer frequency differences and fold changes in CD4^+^ conventional T cells across HLA class II risk extremes. Extremes are defined as the maximum and minimum OR (Table S1), respectively. Differences and fold changes are calculated using the estimated frequency for maximum OR as baseline.

**Figure S3:**
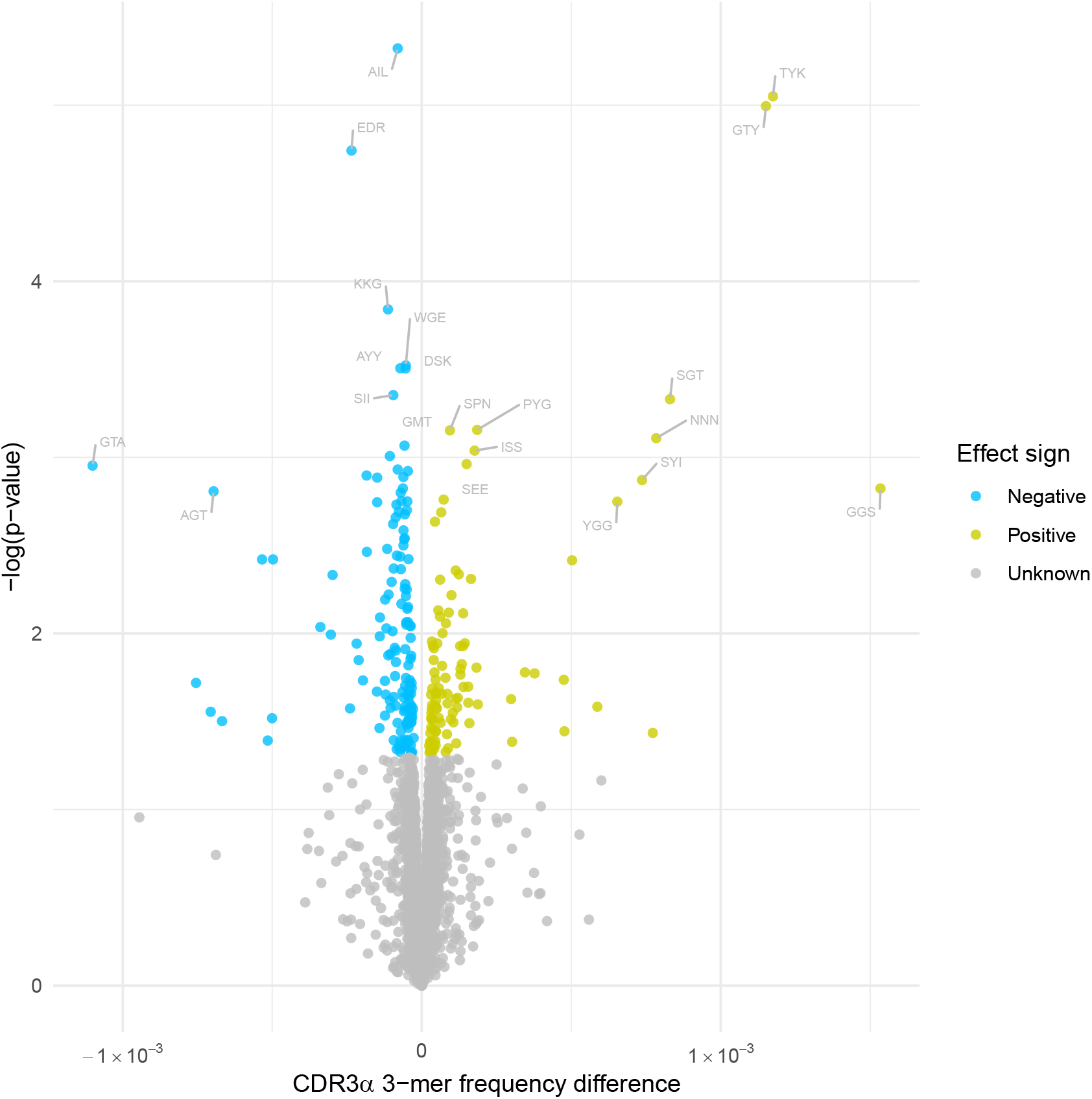
Estimated CDR3α 3-mer frequency differences in CD4^+^ conventional T cells across HLA class II risk extremes. Extremes are defined as the maximum and minimum OR (Table S1), respectively. Differences and fold changes are calculated using the estimated frequency for maximum OR as baseline.

**Figure S4:**
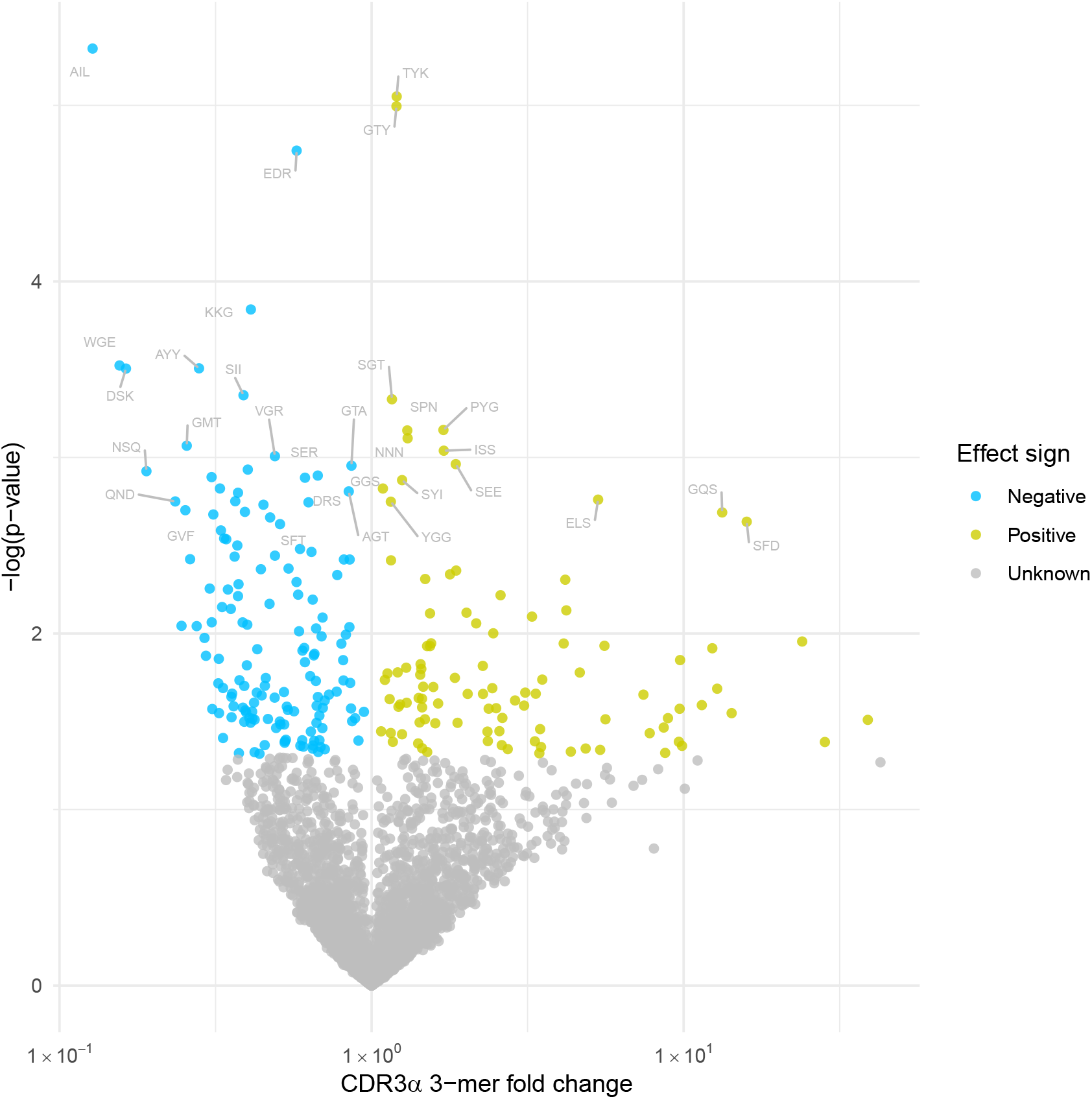
Estimated CDR3α 3-mer fold changes across HLA class II risk extremes. Extremes are defined as the maximum and minimum OR (Table S1), respectively. Differences and fold changes are calculated using the estimated frequency for maximum OR as baseline.

**Figure S5:**
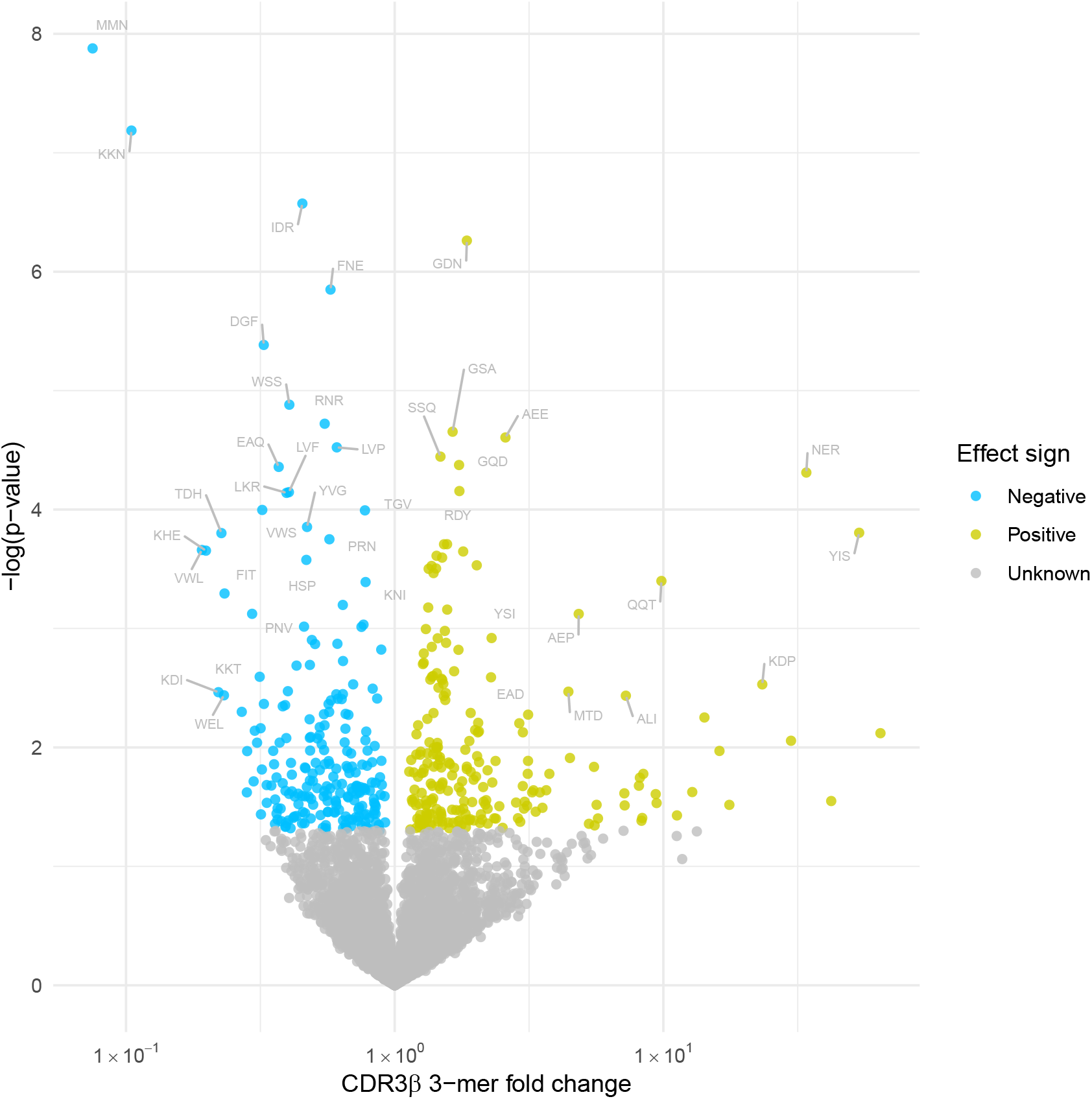
Estimated CDR3β 3-mer fold changes in CD4^+^ conventional T cells across HLA class II risk extremes. Extremes are defined as the maximum and minimum OR (Table S1), respectively. Differences and fold changes are calculated using the estimated frequency for maximum OR as baseline.

**Figure S6:**
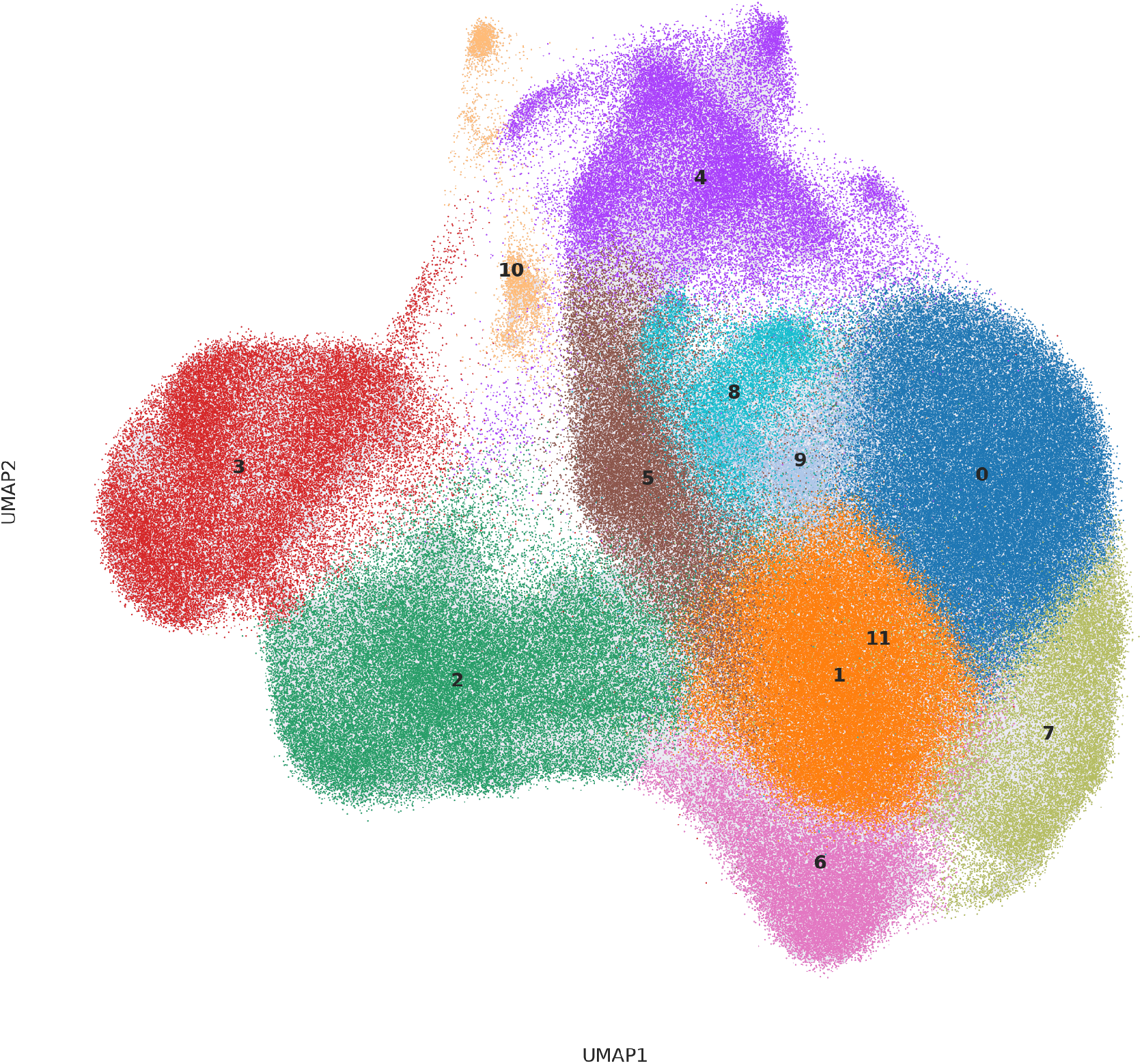
Single-cell RNA clusters. Marker genes for all 12 clusters are depicted next (Figure S7).

**Figure S7:**
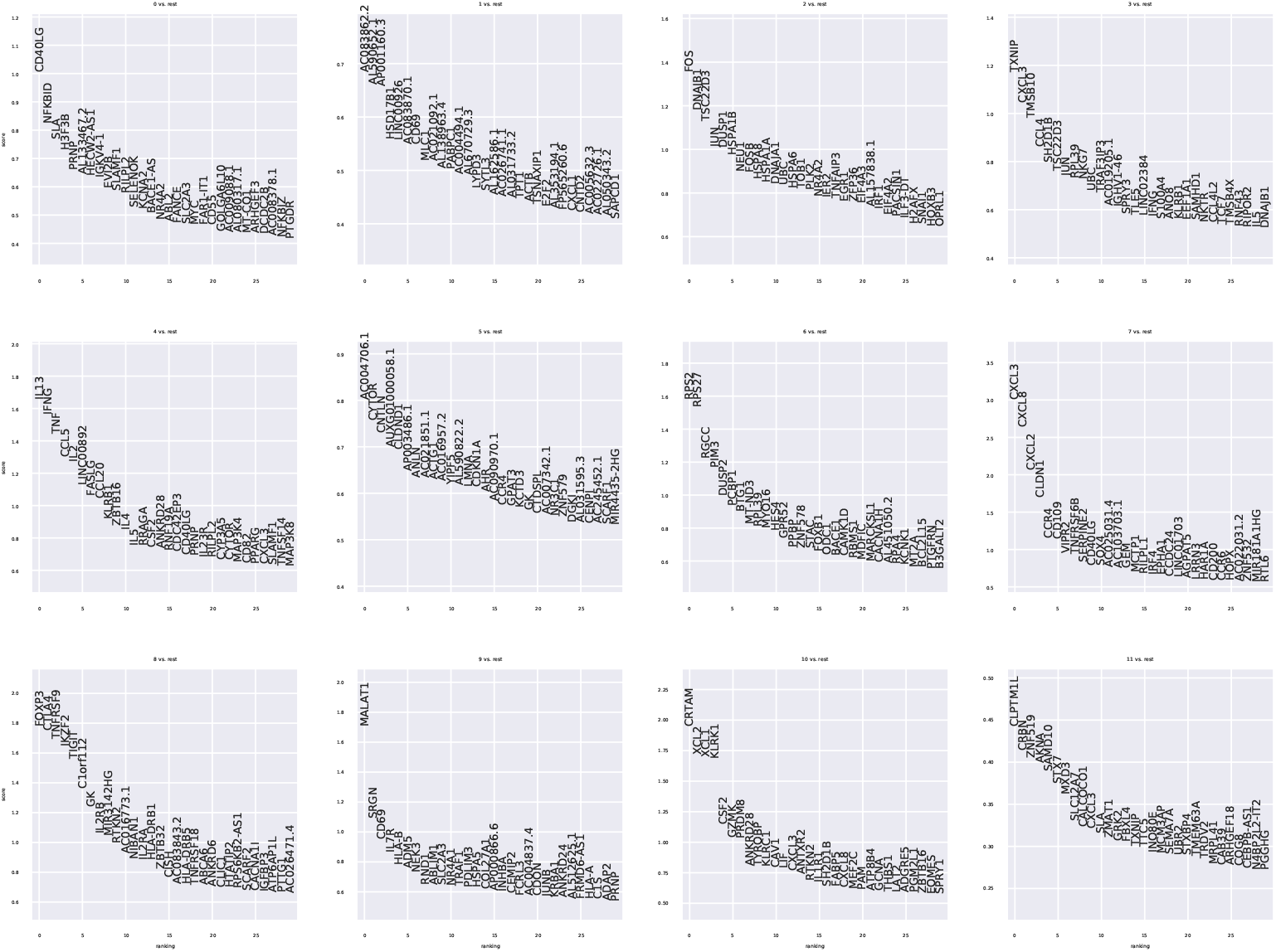
Single-cell RNA marker genes per cluster. Each panel represents a signature consisting of 30 genes that best discriminates [40] a given cluster (Figure S6). Genes are ranked by their score or importance for optimal discrimination.

**Figure S8:**
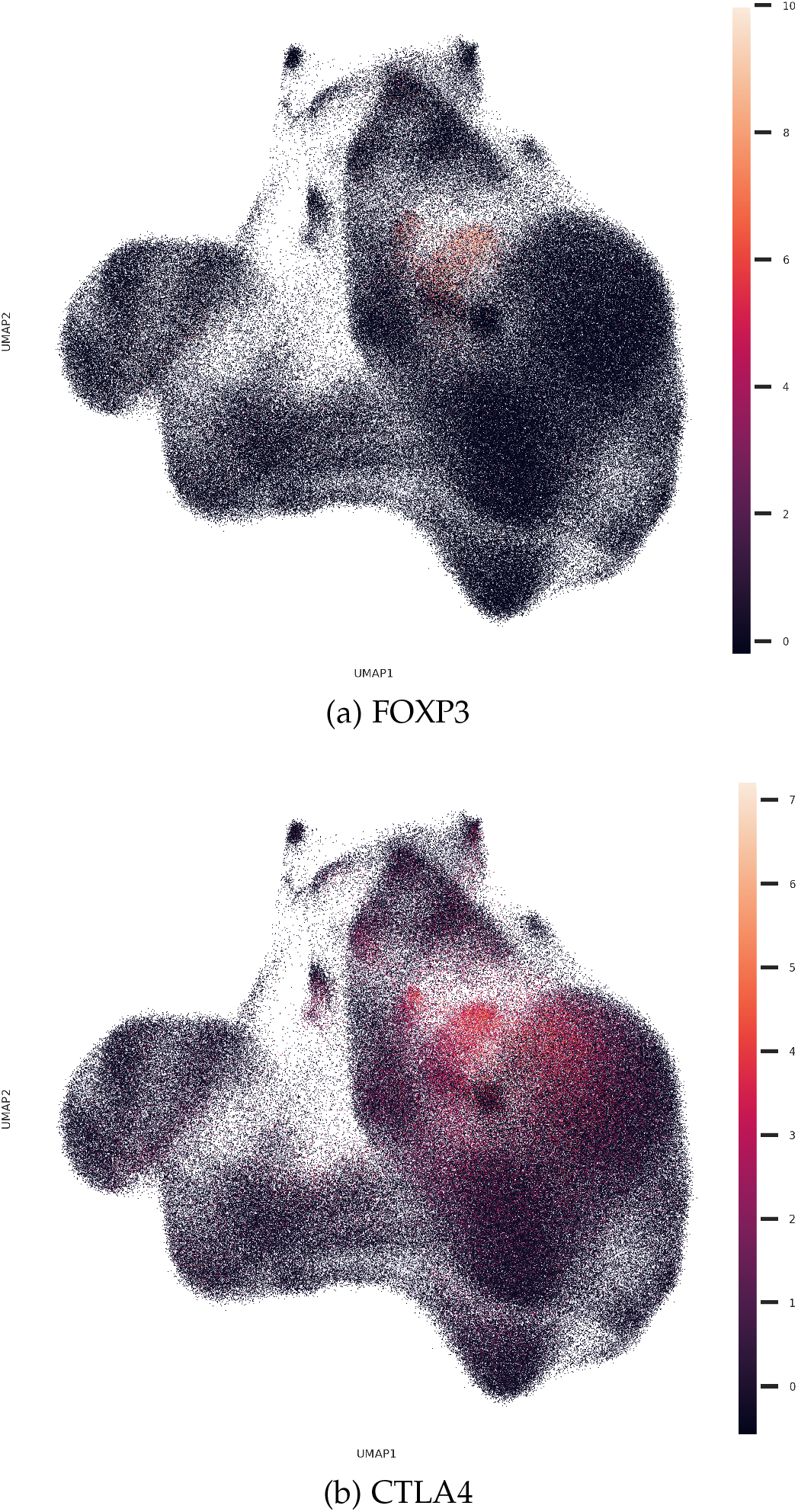
Single-cell RNA regulatory T cell marker expression. Joint expression of classical CD4^+^ Treg markers FOXP3 and CTLA4 maps to predicted Treg cluster (Figures S6 and S7).

## References

[1] American Diabetes Association et al. “2. Classification and diagnosis of diabetes: Standards of Medical Care in Diabetes—2021”. In: Diabetes Care 44.S1 (2021), S15–S33.

[2] John A Todd, John I Bell, and Hugh O McDevitt. “HLA-DQ β gene contributes to susceptibility and resistance to insulin-dependent diabetes mellitus”. In: Nature 329.6140 (1987), pp. 599–604.

[3] Xinli Hu et al. “Additive and interaction effects at three amino acid positions in HLA-DQ and HLA-DR molecules drive type 1 diabetes risk”. In: Nature Genetics 47.8 (2015), p. 898.

[4] Tobias L Lenz et al. “Widespread non-additive and interaction effects within HLA loci modulate the risk of autoimmune diseases”. In: Nature Genetics 47.9 (2015), pp. 1085–1090.

[5] Seth A Sharp et al. “Development and standardization of an improved type 1 diabetes genetic risk score for use in newborn screening and incident diagnosis”. In: Diabetes Care 42.2 (2019), pp. 200–207.

[6] Louis Gioia et al. “Position β57 of I-A^g7^ controls early anti-insulin responses in NOD mice, linking an MHC susceptibility allele to type 1 diabetes onset”. In: Science Immunology 4.38 (2019).

[7] Anette-Gabriele Ziegler et al. “Oral insulin therapy for primary prevention of type 1 diabetes in infants with high genetic risk: the GPPAD-POInT (global platform for the prevention of autoimmune diabetes primary oral insulin trial) study protocol”. In: BMJ Open 9.6 (2019), e028578.

[8] J L Amiel. “Study of the leukocyte phenotypes in Hodgkin’s disease”. In: Histocompatibility Testing. Ed. by E S Curtoni, P L Mattiuz, and R M Tosi. Copenhagen: Munksgaard, 1967, pp. 79–81.

[9] DP Singal and MA Blajchman. “Histocompatibility (HL-A) antigens, lymphocytotoxic antibodies and tissue antibodies in patients with diabetes mellitus”. In: Diabetes 22.6 (1973), pp. 429–432.

[10] J Nerup et al. “HL-A antigens and diabetes mellitus”. In: The Lancet 304.7885 (1974), pp. 864–866.

[11] M Thomsen et al. “MLC typing in juvenile diabetes mellitus and idiopathic Addison’s disease”. In: Immunological Reviews 22.1 (1975), pp. 125–147.

[12] P Platz et al. “HLA-D and -DR antigens in genetic analysis of insulin dependent diabetes mellitus”. In: Diabetologia 21.2 (1981), pp. 108–115.

[13] Arne Svejgaard, Per Platz, and Lars P Ryder. “HLA and disease 1982–a survey.” In: Immunological reviews 70 (1983), pp. 193–218.

[14] Penelope A Morel et al. “Aspartic acid at position 57 of the HLA-DQ beta chain protects against type I diabetes: a family study”. In: Proceedings of the National Academy of Sciences 85.21 (1988), pp. 8111–8115.

[15] K Christopher García et al. “The molecular basis of TCR germline bias for MHC is surprisingly simple”. In: Nature Immunology 10.2 (2009), pp. 143–147.

[16] Eilon Sharon et al. “Genetic variation in MHC proteins is associated with T cell receptor expression biases”. In: Nature Genetics 48.9 (2016), pp. 995–1002.

[17] William S DeWitt III et al. “Human T cell receptor occurrence patterns encode immune history, genetic background, and receptor specificity”. In: eLife 7 (2018), e38358.

[18] Sefina Arif et al. “Blood and islet phenotypes indicate immunological heterogeneity in type 1 diabetes”. In: Diabetes 63.11 (2014), pp. 3835–3845.

[19] Joy A Pai and Ansuman T Satpathy. “High-throughput and single-cell T cell receptor sequencing technologies”. In: Nature Methods (2021), pp. 1–12.

[20] Jacob Glanville et al. “Identifying specificity groups in the T cell receptor repertoire”. In: Nature 547.7661 (2017), pp. 94–98.

[21] Sanzo Miyazawa and Robert L Jernigan. “Residue-residue potentials with a favorable contact pair term and an unfavorable high packing density term, for simulation and threading”. In: Journal of Molecular Biology 256.3 (1996), pp. 623–644.

[22] Stephanie K Lathrop et al. “Peripheral education of the immune system by colonic commensal microbiota”. In: Nature 478.7368 (2011), pp. 250–254.

[23] Dale R Wegmann, Mary Norbury-Glaseru, and Dylan Danielf. “Insulin-specific T cells are a predominant component of islet infiltrates in pre-diabetic NOD mice”. In: European Journal of Immunology 24.8 (1994), pp. 1853–1857.

[24] Maki Nakayama et al. “Prime role for an insulin epitope in the development of type 1 diabetes in NOD mice”. In: Nature 435.7039 (2005), pp. 220–223.

[25] Eddie A James et al. “T-cell epitopes and neo-epitopes in type 1 diabetes: a comprehensive update and reappraisal”. In: Diabetes 69.7 (2020), pp. 1311–1335.

[26] Martha Campbell-Thompson et al. “Network for Pancreatic Organ Donors with Diabetes (nPOD): developing a tissue biobank for type 1 diabetes”. In: Diabetes/Metabolism Research and Reviews 28.7 (2012), pp. 608–617.

[27] Laurie G Landry et al. “Proinsulin-reactive CD4 T cells in the islets of type 1 diabetes organ donors”. In: Frontiers in Endocrinology 12 (2021), p. 217.

[28] Vipin Kumar et al. “The T-cell receptor repertoire and autoimmune diseases”. In: Annual Review of Immunology 7.1 (1989), pp. 657–682.

[29] Eva-Pia Reich et al. “An explanation for the protective effect of the MHC class III–E molecule in murine diabetes”. In: Nature 341.6240 (1989), pp. 326–328.

[30] Howard R Seay et al. “Tissue distribution and clonal diversity of the T and B cell repertoire in type 1 diabetes”. In: JCI Insight 1.20 (2016).

[31] Aaron W Michels et al. “Islet-derived CD4 T cells targeting proinsulin in human autoimmune diabetes”. In: Diabetes 66.3 (2017), pp. 722–734.

[32] Iria Gómez-Touriño et al. “T cell receptor β-chams display abnormal shortening and repertoire sharing in type 1 diabetes”. In: Nature Communications 8.1 (2017), pp. 1–15.

[33] Braden T Tierney et al. “The predictive power of the microbiome exceeds that of genomewide association studies in the discrimination of complex human disease”. In: bioRxiv (2020), pp. 2019–12.

[34] Arcadio Rubio García et al. “Peripheral tolerance to insulin is encoded by mimicry in the microbiome”. In: bioRxiv 2019.12.18.881433 (2019).

[35] Daniel F Zegarra-Ruiz et al. “Thymic development of gut-microbiota-specific T cells”. In: Nature 594.7863 (2021), pp. 413–417.

[36] Xiuwen Zheng et al. “HIBAG—HLA genotype imputation with attribute bagging”. In: The Pharmacogenomics Journal 14.2 (2014), pp. 192–200.

[37] Krzysztof Polański et al. “BBKNN: fast batch alignment of single cell transcriptomes”. In: Bioinformatics 36.3 (2020), pp. 964–965.

[38] Etienne Becht et al. “Dimensionality reduction for visualizing single-cell data using UMAP”. In: Nature Biotechnology 37.1 (2019), pp. 38–44.

[39] Vincent A Traag, Ludo Waltman, and Nees Jan Van Eck. “From Louvain to Leiden: guaranteeing well-connected communities”. In: Scientific Reports 9.1 (2019), pp. 1–12.

[40] Vasilis Ntranos et al. “A discriminative learning approach to differential expression analysis for single-cell RNA-seq”. In: Nature Methods 16.2 (2019), pp. 163–166.

[41] Zachary Sethna et al. “OLGA: fast computation of generation probabilities of B-and T-cell receptor amino acid sequences and motifs”. In: Bioinformatics 35.17 (2019), pp. 2974–2981.

